# Paraspeckles translate microbial insult-induced inflammation into neurovascular remodeling by enhancing CYR61-FGF2 signaling via RBM14 sequestration

**DOI:** 10.64898/2026.07.09.737621

**Authors:** Jia-Qi Pan, Kai-Tong Yang, Jing-Qian Zhang, Yun-Yun Jin, Jian-Huan Chen

## Abstract

**Background:** Systemic inflammation triggered by microbial insults can disrupt endothelial homeostasis, impair blood-brain and blood-retinal barriers, leading to neurovascular remodeling in the central nervous system (CNS). Subnuclear condensates, paraspeckles, play a substantial role in stress-induced gene regulation, yet their contribution to the inflammatory relay from microbial insults to neurovascular remodeling remains unelucidated.

**Results:** Our comparative transcriptomic analysis followed by experimental validation identified a cross-species *NEAT1_2*/CYR61/FGF2 signature in the CNS positively associated with neurovascular remodeling across human disease cohorts and multiple mouse models. Notably, systemic inflammation triggered by microbial insults, including sepsis or gut dysbiosis, enhanced *NEAT1_2* expression in the brain and retina with neurovascular remodeling. Microbial insults induced hyper-assembly of paraspeckles and the expression of CYR61 and FGF2 in vascular endothelial cells. Paraspeckle assembly and its required *NEAT1_2* Domain C, rather than *NEAT1_2* expression levels, play a pivotal role in endothelial homeostasis control and neurovascular remodeling by sequestering the RNA-binding protein RBM14 from the CYR61 promoter, thereby relieving its repression of CYR61 transcription. Moreover, secreted CYR61 enhanced FGF2-mediated endothelial remodeling signals in a paracrine manner. Disrupting paraspeckle assembly by targeting Domain C intercepts neurovascular remodeling, restoring endothelial homeostasis in vivo.

**Conclusions:** Our results demonstrate an essential and conserved role for paraspeckles in the inflammatory relay from microbial insults to neurovascular remodeling by sequestering RBM14 to enhance CYR61-FGF2 signaling. Furthermore, our study underscores paraspeckle assembly as a promising therapeutic target for neurovascular remodeling and related diseases.

## Introduction

Endothelial homeostasis is strictly maintained by quiescence and barrier integrity across the central nervous system (CNS) ^1,2^. Microbial insults, such as infection in sepsis and microbiota dysbiosis, can initiate a systemic-to-local inflammatory relay that allows pro-inflammatory mediators, such as lipopolysaccharide (LPS) endotoxin derived from Gram-negative bacteria, to breach the blood-brain and blood-retinal barriers and disrupt endothelial homeostasis in the CNS ^3–6^. Compromised endothelial homeostasis could, in turn, lead to neurovascular remodeling, including endothelial activation, extracellular matrix remodeling, and pathological neovascularization ^7–10^, the underlying mechanisms of which remain to be further elucidated.

Paraspeckles are membrane-less subnuclear bodies assembled from the long form of long non-coding RNA *NEAT1* (*NEAT1_2*) and paraspeckle proteins ^11–14^. *NEAT1_2*, which harbors Domain C, a key component of paraspeckle assembly, acts as the essential scaffold for this process (**Suppl. Fig. S1**) ^15^. Paraspeckles are known to act as dynamic nuclear hubs that sequester or release RNA-binding proteins (RBPs) in response to cellular stress ^16^ and are implicated in cancer progression and viral infection ^17^. Their roles in endothelial homeostasis and in the systemic-to-local inflammatory relay during neurovascular remodeling remain entirely uncharacterized.

In the current study, we demonstrate a substantial role of paraspeckles in the systemic-to-local inflammatory relay from microbial insults to neurovascular remodeling. Our results showed that microbial insults, stemming from sepsis or gut dysbiosis, trigger inflammation and *NEAT1_2* upregulation. *NEAT1*_2, in turn, promotes paraspeckle hyper-assembly to sequester the RNA-binding protein RNA Binding Motif Protein 14 (RBM14), relieving its transcriptional repression of the pro-angiogenic factor Cysteine-rich angiogenic inducer 61 (CYR61, also known as CCN1). Secreted CYR61 acts in a paracrine manner to mediate endothelial interactions and promotes Fibroblast growth factor 2 (FGF2)-driven remodeling signaling. We also showed that disrupting paraspeckle assembly by targeting *NEAT1_2* Domain C using antisense oligonucleotides (ASOs) efficiently intercepts neurovascular remodeling and restores endothelial homeostasis in vivo. Our findings link microbial insult-triggered inflammation to pathological neurovascular remodeling via a paraspeckle-dependent mechanism, suggesting that paraspeckle integrity is a promising therapeutic target for neurovascular remodeling and related diseases.

## Materials and methods

### Animals

Approval for all animal experiments in the current study was obtained from the Animal Care and Use Committee of Jiangnan University (No. JN.20240530c0700801). 6∼8-week-old C57BL/6J mice were purchased from JOINN Laboratories (Jiangsu, China) and maintained at 20–26°C, 40–70% humidity, with a 12 h light/dark cycle.

### Photocoagulation and neovascularization measurement

Laser-induced choroidal neovascularization (CNV) was used as an in vivo model of neurovascular remodeling in our current study. As previously described ^18^, anesthetized animals received laser photocoagulation at the Bruch’s membrane (532 nm, 0.1 s, 120 mW) and were sacrificed on Day 3 post-laser induction, a time point with the most compelling transcriptomic changes in the retina ^18^. Flat mounts of the retina and choroid were performed and stained with FITC-Isolectin B4 (Enzo Life Sciences, ALX-650-001F-MC05, USA), then imaged using a Zeiss LSM800 confocal microscope, and CNV lesion was quantified using ImageJ.

### Intravitreal injection

Intravitreal injections in mice were performed via a trans-scleral approach with a 33-gauge Hamilton syringe.

### Gut dysbiosis and antibiotic depletion models

Mice were gavaged daily with Escherichia coli (E. coli) (2 × 10⁸ CFU in 200 μL) for 7 days. Laser photocoagulation was performed on day 4, and mice were sacrificed on day 7. Cecal contents were collected for 16S rRNA sequencing; colon tissues were processed for H&E staining. For microbiota depletion, mice received a broad-spectrum antibiotic cocktail (ampicillin 3 mg/mL, neomycin 3 mg/mL, metronidazole 3 mg/mL, vancomycin 1.5 mg/mL in PBS; 300 μL every other day for 7 days) starting before laser treatment and continuing until endpoint.

### 16S rRNA sequencing and data analysis

Illumina HiSeq sequencing of cecal microbiota was performed by Novogene (China). Raw data were analyzed using QIIME2 (v2020.11) ^19^. α- and β-diversity (weighted UniFrac) were calculated, and differentially abundant taxa were then identified using linear discriminant analysis (LDA) Effect Size (LEfSe) method (LDA score > 2) ^20^.

### Plasmids, siRNA, and ASOs

Full-length Human *CYR61* and *RBM14* cDNAs were cloned into pcDNA3.1(+) via PCR from HEK293T cells. The *CYR61* promoter was amplified from genomic DNA and cloned into pGL3-Basic (pGL3-CYR61). A deletion mutant with a predicted RBM14 binding site was constructed using a mutagenesis kit (Vazyme, C216, China). ASOs and siRNA were purchased from GenePham (Suzhou, China). Sequences of Primer, siRNA, and ASO are provided in **Supplementary Tables 1-2**.

### Cell culture, transfection, and stimulation

Human Retinal Microvascular Endothelial Cells (HRMECs), Human adult retinal pigment epithelial (ARPE)-19 cells, and Human Embryonic Kidney 293T Cells (HEK293T) were cultured in DMEM (Cytiva, SH30022.01, USA) supplemented with 10% heat-inactivated fetal bovine serum (CELLiGENT, CG0430A, New Zealand) at 37°C in 5% CO₂. Lipofectamine 3000 (Invitrogen, L3000015, USA) was used for transfections. HRMECs were stimulated with LPS (Sigma, L2630, USA; 50 μg/mL) for 12 h prior to RNA extraction.

### RNA extraction and qPCR

Total RNA was extracted (TIANGEN, Y1920, China), reverse-transcribed (Vazyme, R323, China), and quantified via qPCR using ChamQ SYBR Master Mix (Vazyme, Q711, China) on a Roche LightCycler 480 II. Primers are in **Supplementary Table 3**.

### RNA sequencing and public dataset retrieval

Bulk and single-nucleus RNA-seq datasets were downloaded from NCBI Gene Expression Omnibus (http://www.ncbi.nlm.nih.gov/geo) and the Single Cell Portal (https://singlecell.broadinstitute.org/single_cell/study/SCP2012). Datasets included in the current study are listed as follows: GSE115828 ^21^ of human retina, AMD cohort (controls and 3 disease stages), n=453; GSE236562 ^22^ of human brain (Brodmann area 9 and hippocampal formation), AD and age/gender-matched controls, n=8; GSE237861 ^23^, human prefrontal cortex, septic shock and non-infectious controls (n=14) and SCP2012 (sNuc-seq) ^24^ of human retina/RPE/choroid with 164,399 cells from 6 AMD and 7 control donors; GSE158799 of the retina from oxygen-induced retinopathy (OIR) mice at P14 (n=2), P17 (n=3), P30 (n=3), and controls (n=2); PRJNA781828 of the retina from NaIO₃-induced degeneration at day 3 (n=5) and control mice (n=5); GSE223522 of the retina from diabetic retinopathy mice at 6 weeks (n=5), 8 weeks (n=5), and 10 weeks (n=5) and controls (n=6).

### RNA-seq data processing and analysis

Raw reads were processed using FASTP version 0.23.4 ^25^ to remove low-quality reads and mapped to the UCSC mouse mm10 or human hg38 reference genome using STAR version 2.7.0e ^26^. Gene-level pseudo-counts were quantified using FeatureCounts version 2.0.1 ^27^ and used for TPM value calculation using edgeR version 3.7 ^28^. Differential expression was assessed at *P* < 0.05. Spearman’s correlation analysis was performed to assess gene co-expression. Enrichment analysis was conducted using Enrichr (https://maayanlab.cloud/Enrichr/) ^29^ and DAVID (https://david.ncifcrf.gov/tools.jsp) ^30^. *NEAT1* isoform expression was analyzed using StringTie version 2.2.1 ^31^, and subjected to normalized counts calculation implemented in DESeq2 version 3.22 ^32^.

### Western blotting

Protein extraction was conducted using the protein extraction kit with 1% protease inhibitor cocktail (CWBIO, CW0043S, China). Concentrations were determined by the BCA assay. Total protein was run on 10% SDS-PAGE and probed with antibodies against CYR61 (ABclonal, A1111, China; 1:1000), FGF2 (Proteintech, 11234-1-AP, China; 1:1000), RBM14 (Proteintech, 10196-1-AP, China; 1:1000), and ACTB (ABclonal, AC026, China; 1:2000).

### Luciferase reporter and endothall functional assays

Luciferase assays were performed in HEK293T cells co-transfected with pGL3-CYR61 constructs and pcDNA3.1-RBM14 using Lipofectamine 3000, followed by Dual-Luciferase Reporter Kit (Beyotime, RG027, China). Cell proliferation was assessed via CCK-8 (Beyotime, C0038, China) on a BioTek Synergy H4 microplate reader. Wound-healing and tube formation assays were performed by following standard protocols. Cell images were captured with a Nikon microscope and quantified using ImageJ.

### Immunofluorescence, RNA-FISH, and ChIP-qPCR

HRMECs were fixed with 4% PFA (Solarbio), permeabilized with 0.1% Triton X-100, blocked with 5% goat serum, and incubated overnight with anti-RBM14 antibody (Boster, A04917-2, China). Cells were stained with Alexa Fluor 647 secondary antibody (Beyotime, A0468, China) and hybridized with Cy3-labeled *NEAT1_2* probes (Ribobio, C10920, China). Nuclei were counterstained with DAPI (Solarbio, S2110, USA) and imaged on a Zeiss LSM800. ChIP was performed using a kit (Beyotime, P2080S, China), anti-RBM14 antibody (Abcam, ab70636, USA), and DNA purified with a Beyotime kit (D0033, China). ChIP-qPCR targeted the CYR61 promoter region (primers in **Supplementary Table 4**).

### Paraspeckle structural disruption and ELISA

HRMECs were treated with 10% 1,6-hexanediol (1,6-HD) or 10% 2,5-hexanediol (2,5-HD) for 30 min ^33^. CYR61 secretion and plasma LPS levels were measured using Elabscience kits (E-EL-H1285 and E-EL-0180, China).

### Statistical analysis

Data are shown as mean ± SD with ≥ 3 independent triplicates. Statistical difference was analyzed using Student’s t-test for two-group comparison or one-way ANOVA for comparison of three or more groups using SPSS 27.0. *P* < 0.05 was considered significant.

## Results

### Cross-species comparative transcriptomics identifies a conserved upregulated NEAT1_2/CYR61/FGF2 signature associated with neurovascular remodeling

To examine the transcriptomic changes underlying neurovascular remodeling, we performed a high-resolution cross-species transcriptomic analysis integrating RNA-seq data from human cohorts of AMD (GSE115828) and AD (GSE236562) (**Fig. 1A**), as well as neurovascular remodeling mouse models, including laser-induced CNV for exudative AMD from our current study, OIR for retinopathy (GSE158799), NaIO₃-induced degeneration for dry AMD (PRJNA781828), and DR (GSE223522). Among differentially expressed lncRNAs across these pathological contexts, *NEAT1* and its isoform *NEAT1_2* showed consistent upregulation across both human cohorts (**Fig. 1A, Suppl. Fig. S2**) and four mouse models (**Fig. 1B, C**), and were validated in the CNV mouse retina by qPCR (**Fig. 1D**).

**Figure 1.**
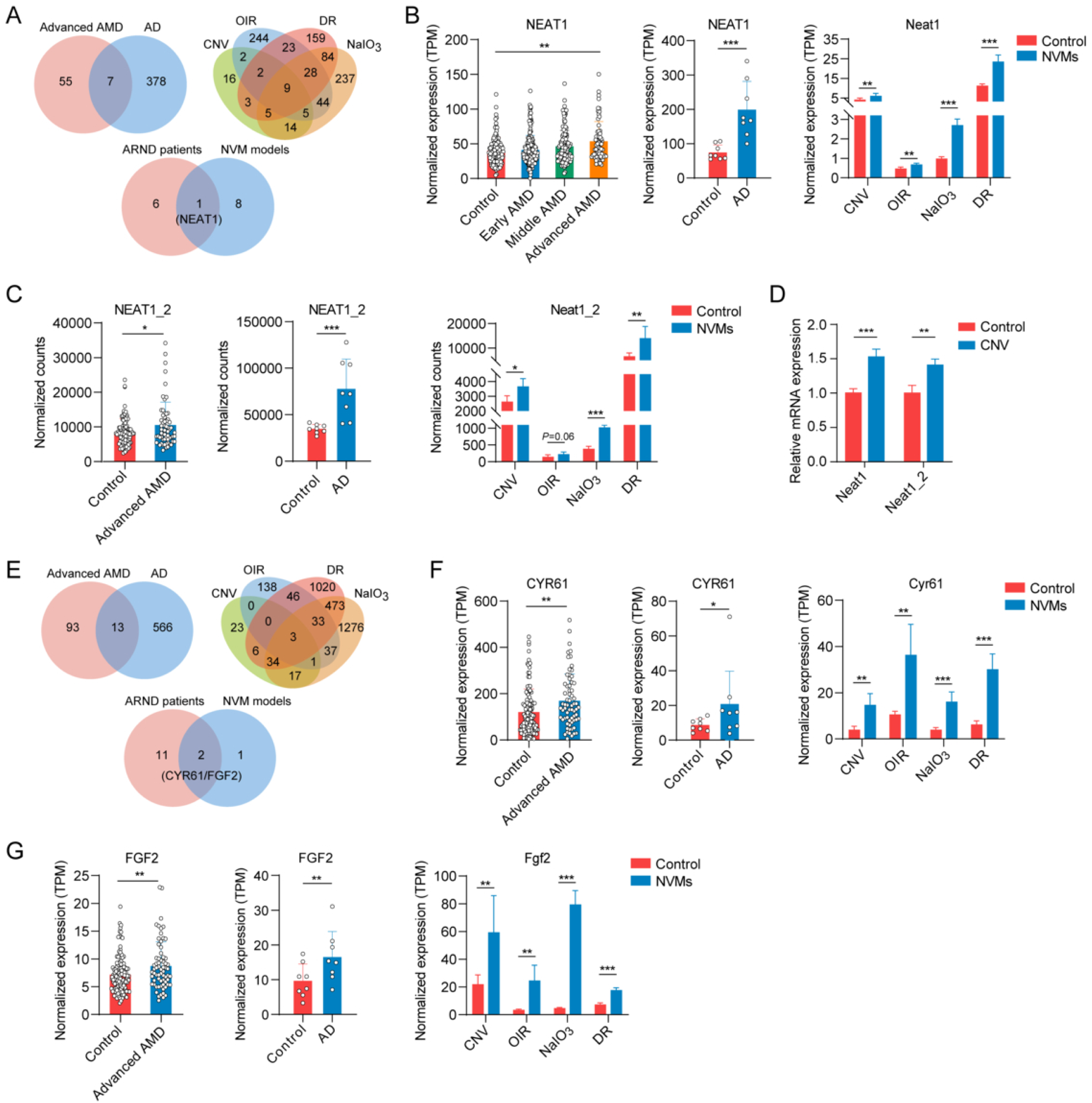
Cross-species transcriptomic profiling reveals a conserved upregulated *NEAT1_2*/CYR61/FGF2 signature across human cohorts and mouse models of neurovascular remodeling. (**A**) Venn diagram comparing differentially expressed lncRNAs between human AMD and AD cohorts and multiple mouse models of neurovascular remodeling. (**B, C**) Normalized expression of *NEAT1* and its long isoform *NEAT1_2* across stages of human cohorts of AMD and AD, and neurovascular remodeling mouse models. (**D**) Validation of *NEAT1/NEAT1_2* in the CNV retina using qPCR. (**E**) Venn diagram identifying *NEAT1*-co-expressed DEGs overlapping between human ARNDs and murine NVP models (Spearman |*r*| ≥ 0.5, *p* < 0.05). (**F, G**) Normalized expression of *CYR61* and *FGF2* in human neurodegenerative disease cohorts and neurovascular remodeling-related mouse models. Data are shown as mean ± SD from ≥ 3 independent triplicates. \**p* < 0.05, \*\**p* < 0.01, \*\*\**p* < 0.001. NVM: neurovascular remodeling

To explore the regulatory networks associated with *NEAT1*, we performed a co-expression analysis of cross-species comparative transcriptomic data. Angiogenic factors, CYR61 and FGF2, were identified as the only two *NEAT1*-coexpressed DEGs exclusively across the human cohorts and murine models analyzed above (Spearman *r* = 0.5, *p* < 0.05; **Fig. 1E, Suppl. Fig. S3**), both positively correlated with *NEAT1_2* (**Fig. 1F, G, Suppl. Fig. S4A-F**), and further validated in the CNV mouse retina (**Suppl. Fig. S4G**). Such findings thus pointed to a conserved upregulated *NEAT1_2*/CYR61/FGF2 signature associated with neurovascular remodeling.

### Inflammation triggered by microbial insults induced NEAT1_2 upregulation, leading to paraspeckle hyper-assembly

Our subsequent results showed higher *NEAT1* and *NEAT1_2* expression in the retina of AMD patients with sepsis than in those without sepsis (**Fig. 2A**), and in the prefrontal cortex of septic patients than in controls (GSE237861; **Fig. 2A**), suggesting that microbial insult-induced inflammation may increase *NEAT1* and *NEAT1_2* expression. To validate such findings, we used an *E. coli*-induced dysbiosis model to test whether microbial insult-induced inflammation affects neovascular remodeling in laser-induced CNV mice (**Fig. 2B**). Gut dysbiosis was confirmed by changes in alpha-diversity and beta-diversity (**Fig. 2C, Suppl. Fig. S5A-C**), as well as by colonic inflammation (**Fig. 2D**) and decreased tight-junction gene expression (**Suppl. Fig. S5D**). Gut dysbiosis increased plasma LPS levels (**Fig. 2E**), a hallmark of microbial insult-induced inflammation. LPS enhanced neurovascular remodeling, resulting in more severe CNV lesions and increased astrocyte and microglial activation, as indicated by GFAP⁺ and IBA1⁺ staining, pointing to exacerbated neuroinflammation (**Fig. 2F, G, and Suppl. Fig. S5E**). The LPS sensing pathway (Tlr4, Myd88, Il6st; **Suppl. Fig. S5F**) and the *NEAT1_2*/CYR61/FGF2 signature were upregulated in the CNV mouse retina (**Fig. 2H, I**). In contrast, antibiotic treatment suppressed CNV, confirming that gut dysbiosis-induced systemic inflammation drove neurovascular remodeling (**Fig. 2J**).

**Figure 2.**
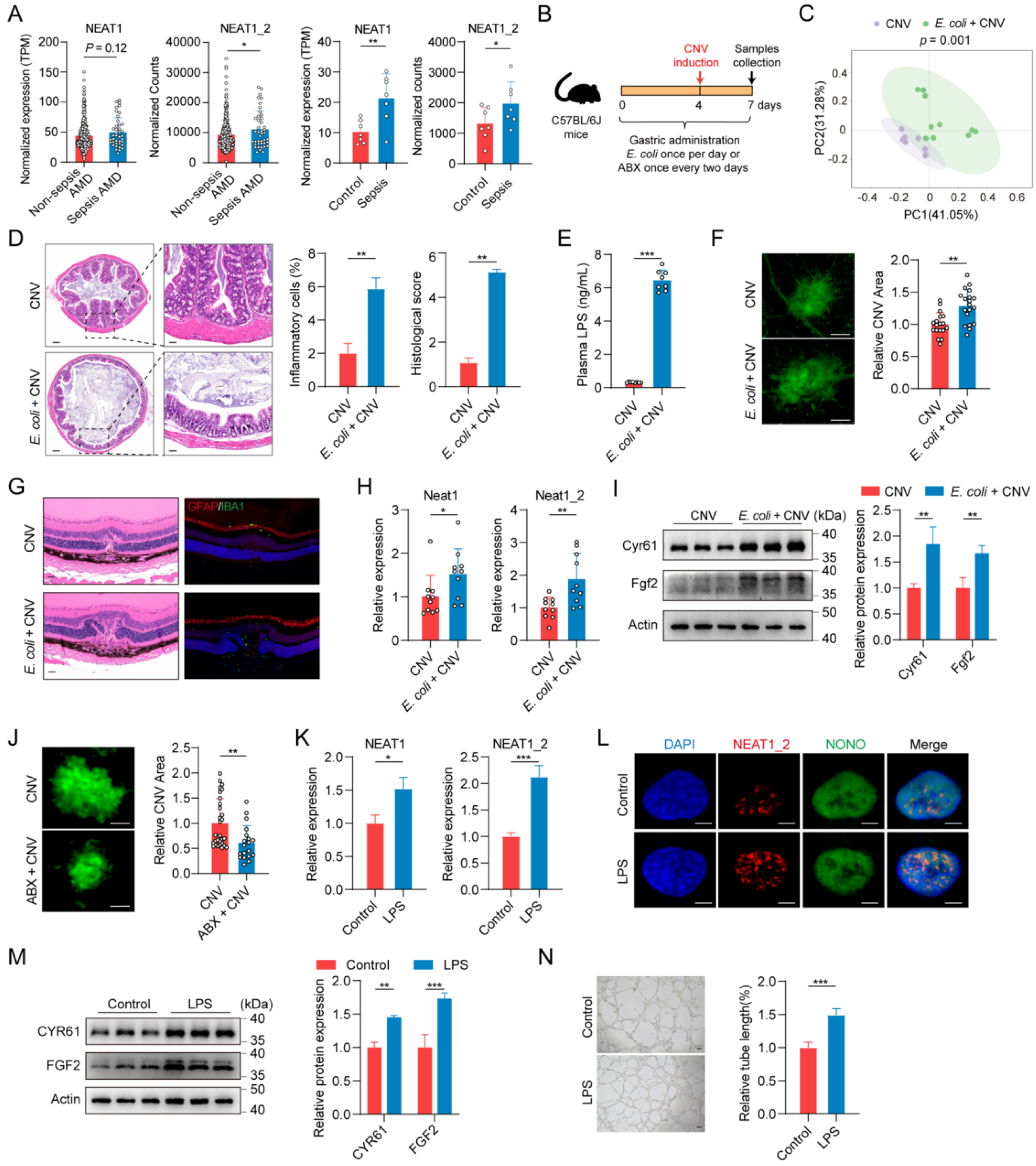
Systemic inflammation triggers neurovascular remodeling and upregulation of *NEAT1_2*, associated with paraspeckles hyper-assembly. (**A**) Clinical cohort analysis of *NEAT1/NEAT1_2* expression in AMD patients with or without sepsis, and in the prefrontal cortex of septic individuals. (**B**) Diagram showing experimental workflow of *E. coli*-induced gut dysbiosis and antibiotic cocktail (ABX)-mediated microbiota depletion in CNV mice. (**C**) PCoA of β-diversity derived from 16S rRNA sequencing of gut microbiota following *E. coli* treatment; *n* = 8–10 per group). (**D**) H&E staining of colonic tissue and the quantification of inflammatory infiltration and histological scores. Scale bars: left, 50 μm; right, 200 μm. (**E**) Plasma LPS levels measured using ELISA (*n* = 8 per group). (**F**) Isolectin-B4-stained choroidal flat mounts showing CNV. Scale bar = 100 μm. (**G**) H&E (left) and immunofluorescence staining (right) in retinal section. GFAP (red) and IBA1 (green), and DAPI (blue) for Nuclei are stained. Scale bar = 200 μm. (**H, I**) qPCR and western blot of *NEAT1/Cyr61/Fgf2* RNA and protein upregulation in the retina (*n* = 10 for qPCR and 5 for WB per group). (**J**) Isolectin-B4 staining demonstrating ABX-mediated suppression of CNV progression. (**K, L**) Confocal microscopy showing LPS-induced *NEAT1_2*/NONO colocalization indicative of paraspeckle hyper-assembly in HRMECs. Scale bar = 5 μm. (**M**) Western blot of CYR61 and FGF2 upon LPS stimulation. (**N**) Tube formation assays post-LPS treatment (12 hrs). Data are shown as mean ± SD from ≥ 3 independent triplicate measurements. \**p* < 0.05, \*\**p* < 0.01, \*\*\**p* < 0.001.

Furthermore, our in vitro experiments showed that inflammation in HRMECs induced by LPS upregulated expression of *NEAT1* and *NEAT1_2* and promoted paraspeckle assembly, as indicated by *NEAT1_2* colocalizing with paraspeckle component Non-POU domain-containing octamer-binding protein (NONO) (**Fig. 2K, L**), with upregulated pro-angiogenic factors CYR61 and FGF2, as well as enhanced endothelial proliferation, migration, and tube formation (**Fig. 2M, N** and **Suppl. Fig. S5G-J**).

These findings showed that microbial insult-induced inflammation induced by sepsis and gut dysbiosis could drive neurovascular remodeling through hyper-assembly of *NEAT1_2*-scaffolded paraspeckles.

### NEAT1_2 Domain C and paraspeckle assembly are required for vascular endothelial homeostasis regulation

To determine the role of *NEAT1_2* and paraspeckle assembly in vascular endothelial homeostasis regulation, we used ASOs to identify the functional domains of *NEAT1_2* in HRMECs, including **Domain A** responsible for stabilization, **Domain B** for isoform switching, and **Domain C** for paraspeckle assembly (**Fig. 3A**). CCK-8, wound-healing, and tube-formation assays showed all three ASOs inhibited activity, but disrupting Domain C showed the strongest suppressing effects (**Fig. 3B, Suppl. Fig. S6A, B**). These findings suggest that paraspeckle assembly, rather than the presence of NEAT1 alone, is more important for endothelial activation. Moreover, qPCR and western blot showed that Domain C knockdown led to the strongest decrease in CYR61 expression (**Fig. 3C, Suppl. Fig. S6C**). Consistently, siRNA knockdown of CYR61 inhibited proliferation, migration, and angiogenesis in HRMECs, consistent with the *NEAT1_2* Domain C disruption results (**Fig. 3D, Suppl. Fig. S6D-F**). These findings thus suggest a causal link between subnuclear architectural integrity and vascular endothelial remodeling. These findings indicate that Domain C is essential for paraspeckle hyper-assembly, subsequent CYR61 induction, and regulation of vascular endothelial homeostasis.

**Figure 3.**
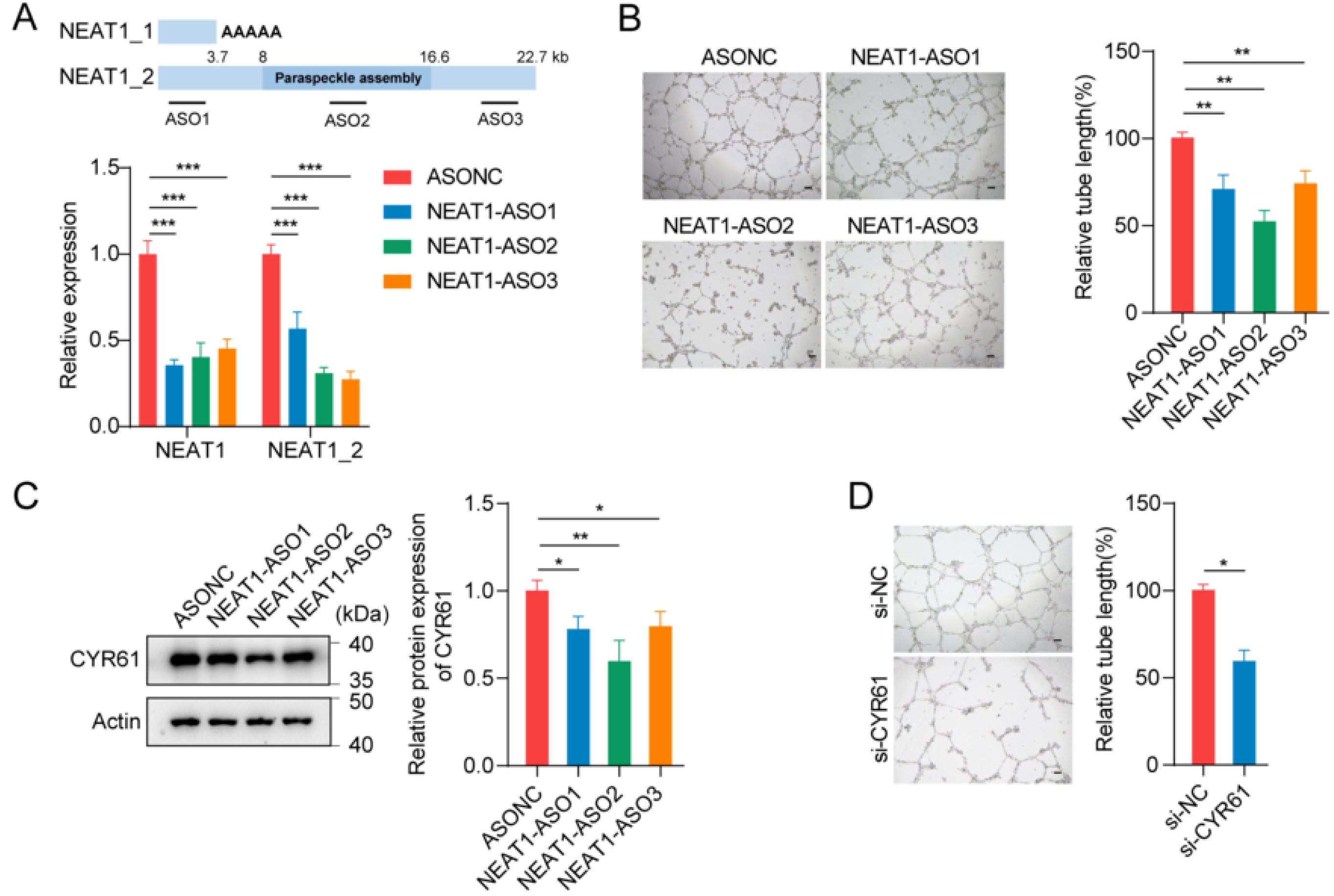
Structural integrity of paraspeckles is required for CYR61 activation and pro-angiogenic signaling in vascular endothelium. (A) Schematic of ASOs targeting domains responsible for stabilization (Domain A), isoform switching (Domain B), and paraspeckle assembly (Domain C), with qPCR validation of knockdown efficiency. (B) Tube formation assays in HRMECs transfected with *NEAT1* ASOs. (C) Western blot analysis of CYR61 expression following domain-specific ASO treatment. (D) Tube formation assays in HRMECs with CYR61 knockdown. Data are shown as mean ± SD from ≥3 independent triplicates. Scale bar = 100 μm. \**p* < 0.05, ** *p* < 0.01, *** *p* < 0.001.

### Paraspeckle assembly sequesters RBM14, relieving the transcriptional brake on CYR61

To further identify regulatory factors involved in CYR61 expression regulation by paraspeckle assembly, we used GTRD to analyze core paraspeckle proteins and predicted RBM14 and NONO binding sites in the *CYR61* promoter (**Fig. 4A, Suppl. Fig. S7A**). Further functional screening showed that knocking down *RBM14*, not *NONO*, increased CYR61 expression in HRMECs (**Suppl. Fig. S7B, C**). Moreover, disruption of *NEAT1_2* Domain C by ASOs did not alter RBM14 mRNA or protein levels (**Fig. 4B**). *NEAT1_2* ASO2 targeting Domain C disrupted paraspeckle formation, indicated by reduced colocalization of *NEAT1* FISH with RBM14 immunofluorescence and relocated RBM14 into the nucleoplasm (**Fig. 4C**).

**Figure 4.**
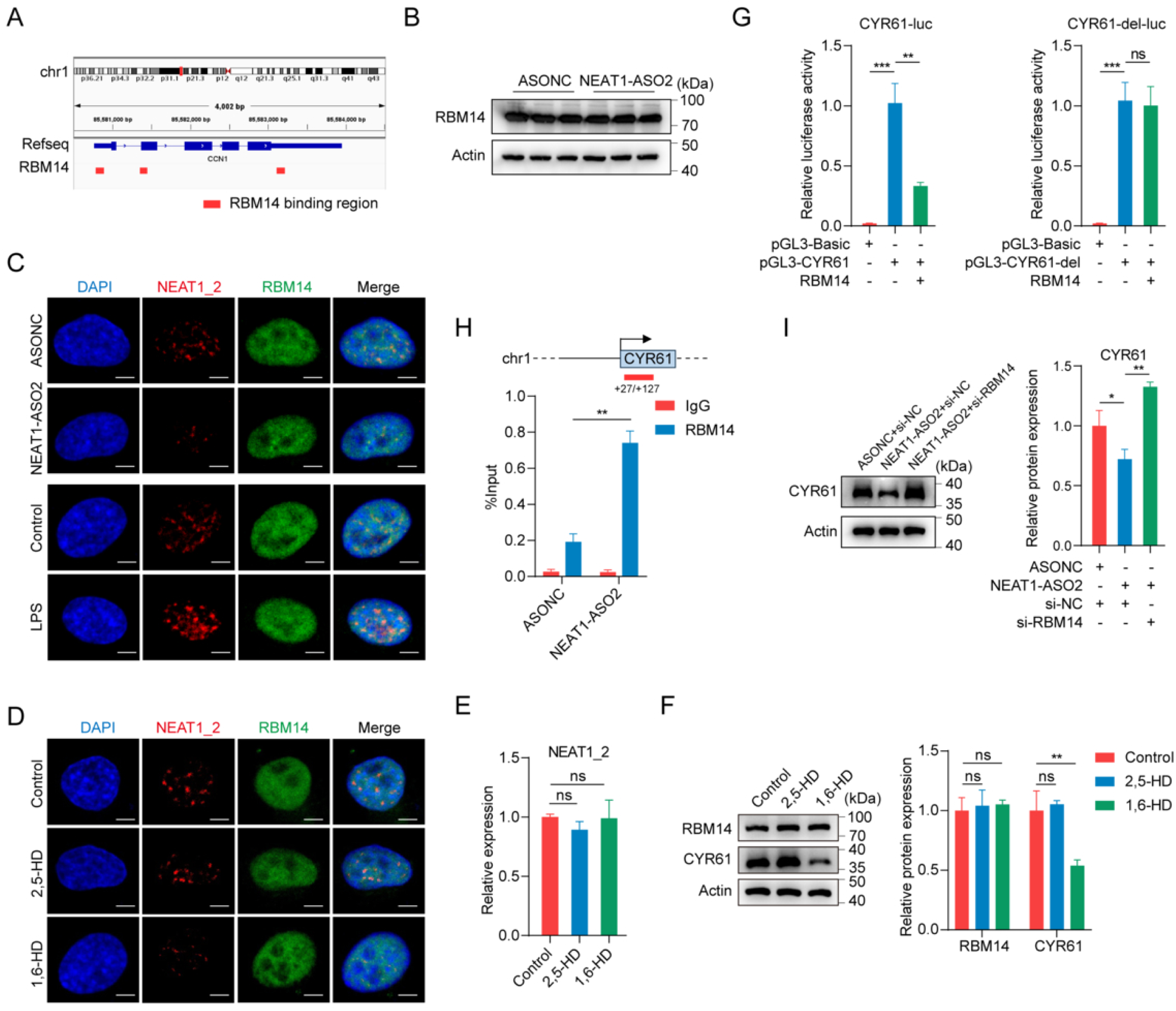
Paraspeckle assembly sequesters RBM14, relieving CYR61 transcriptional repression. (**A**) RBM14 binding sites within the proximal CYR61 promoter predicted by GTRD. (**B, C**) *NEAT1* ASO2-mediated disruption of **paraspeckle assembly** alters subnuclear RBM14 localization without affecting RBM14 protein expression, as confirmed by FISH/IF. (**D-F**) for 30 min-treatment with 10% 1,6-HD, but not the 2,5-HD control, disrupts paraspeckle assembly and downregulates CYR61, with *NEAT1_2*/RBM14 expression unchanged. (**G**) Dual-luciferase assays showing RBM14-mediated repression of wild-type CYR61 promoter activity, which is abolished upon motif deletion (+27 to +127 bp). (**H**) ChIP-qPCR confirming direct RBM14 binding to the *CYR61* promoter, which is enhanced upon *NEAT1_2* knockdown. (**I**) RBM14 knockdown using siRNA rescues *NEAT1* ASO2-induced CYR61 expression suppression. Data are shown as mean ± SD from ≥ 3 independent triplicates. Scale bar = 5 μm. \**p* < 0.05, \*\**p* < 0.01, \*\*\**p* < 0.001, ns: not significant.

These findings were further supported by paraspeckle structural disruption using 1,6-hexanediol (1,6-HD), which disrupts paraspeckles without altering *NEAT1_2* or RBM14 expression levels. (**Fig. 4D-F**). Such findings, together with the Domain C-targeting ASO results, confirmed that paraspeckle assembly regulates CYR61 expression via RBM14 sequestration. Luciferase reporter assays further showed that RBM14 repressed the wild-type *CYR61* promoter, an effect abolished by deletion of the predicted binding site (+27 to +127 bp upstream of the CYR61 transcription start site) **(Fig. 4G)**. RBM14 binding to the CYR61 promoter was confirmed by ChIP-qPCR, and its occupancy increased after *NEAT1_2* ASO2 treatment (**Fig. 4H**), linking loss of sequestration to repression. Consequently, RBM14 knockdown rescued the CYR61 downregulation induced by *NEAT1_2* ASO2 (**Fig. 4I, Suppl. Fig. S7D**), indicating that paraspeckles sequester RBM14 and promote CYR61 transcription, thereby promoting neovascular remodeling.

### Paraspeckle hyper-assembly orchestrates paracrine CYR61/FGF2 signaling and promotes neovascular remodeling

To explore the downstream pathway of CYR61, we conducted RNA-seq on HRMECs overexpressing or knocking down CYR61 (**Suppl. Fig. S8A, B**). Our results identified a set of 559 genes co-expressed with *CYR61* (**Fig. 5A**), mainly involved in “ameboidal-type cell migration” and “tissue remodeling,” consistent with the potential role of CYR61 in endothelial homeostasis (**Fig. 5B**). Further filtering this gene set using the DEGs in human and mouse transcriptomic data in **Fig. 1** identified 10 conserved hub genes, particularly *FGF2*, a key factor in neurovascular survival and angiogenesis (**Fig. 5C, Suppl. Fig. S8C, D**), which was confirmed by both western blot and qPCR (**Fig. 5D-E, Suppl. Fig. S8E**). Overexpressing *CYR61* rescued the reduction of FGF2 expression induced by ASO2-mediated disruption of *NEAT1_2* Domain C, confirming FGF2 as a downstream effector regulated by the *NEAT1_2*-scaffolded paraspeckle assembly (**Fig. 5F-G, Suppl. Fig. S8F-G**).

**Figure 5.**
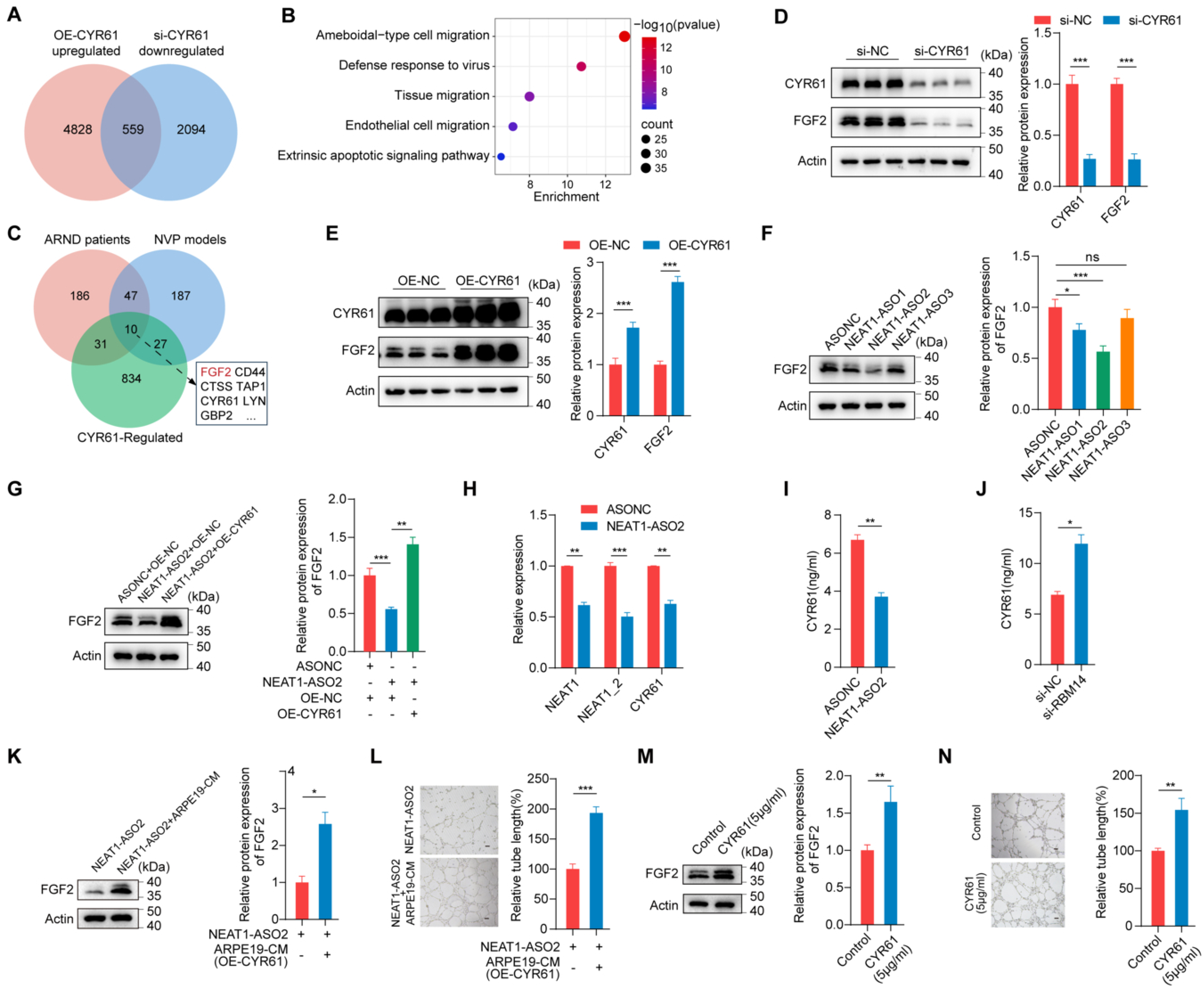
CYR61 paracrine drives FGF2 signaling in endothelial cells and promotes neovascular remodeling. (**A**) Venn diagram showing a set of 559 DEGs positively regulated by CYR61, which are upregulated by CYR61 overexpression and downregulated by CYR61 knockdown. (**B**) Top five GO biological process terms enriched for the DEGs positively regulated by CYR61. (**C**) Intersection of the DEGs positively regulated by CYR61 with DEGs from advanced AMD and CNV mouse models. (**D, E**) Western blot confirming strict dependency of FGF2 expression on the *NEAT1_2*-CYR61 axis. (**F, G**) Domain C disruption downregulates FGF2, an effect significantly rescued by CYR61 overexpression. (H-J) ARPE-19 assays showing that Domain C disruption suppresses CYR61 secretion, whereas RBM14 knockdown enhances CYR61 secretion. (K, L) Conditioned media from ARPE-19 cells overexpressing CYR61 rescues *NEAT1* ASO-induced suppression of HRMEC angiogenesis and FGF2 expression. (**M, N**) Recombinant human CYR61 directly upregulates FGF2 and enhances tube formation in HRMECs. Data are shown as mean ± SD in triplicate. Scale bar = 100 μm. \**p* < 0.05, **p < 0.01, \*\*\**p* < 0.001, ns: not significant.

As single-nucleus RNA-seq data showed widespread expression of *NEAT1_2*, *CYR61*, and *FGF2* across cell types (**Suppl. Fig. S8H**), we then evaluated the potential paracrine effect of CYR61. Regulation of CYR61 secretion through the *NEAT1_2*-RBM14 axis was confirmed in human ARPE19 cells, consistent with findings in HRMECs (**Fig. 5H-J, Suppl. Fig. S8I**). Moreover, culture medium from RPE cells overexpressing CYR61 rescued tube formation and FGF2 expression that were suppressed by either ASO2-mediated *NEAT1_2* Domain C disruption or by CYR61 knockdown in HRMECs. (**Fig. 5K–L, Suppl. Fig. S8J–M**), which was confirmed by increased FGF2 expression and enhanced tube formation in HRMECs supplemented with recombinant CYR61 (**Fig. 5M-N, Suppl. Fig. S8N**). These findings demonstrate that *NEAT1_2*/paraspeckle hyper-assembly orchestrates paracrine CYR61/FGF2 signaling and promotes neovascular remodeling.

### Targeted disruption of NEAT1_2 paraspeckle assembly intercepts neovascular remodeling in vivo

To evaluate the translational potential of modulating subnuclear architecture in vivo, we used a localized therapeutic approach by intravitreal injection of ASOs targeting the *NEAT1_2* paraspeckle assembly Domain C. Isolectin-B4-staining of choroidal flat mounts showed that targeted disruption of Domain C resulted in a marked reduction in CNV **(Fig. 6A)**. *NEAT1_2* ASO treatment also effectively suppressed the expression of Cyr61 and Fgf2 in the retina (**Fig. 6B**). These findings show that targeting Domain C of the *NEAT1_2* blocks neovascular remodeling and restores endothelial homeostasis, suggesting a potential therapeutic approach for neurovascular remodeling and related diseases.

**Figure 6.**
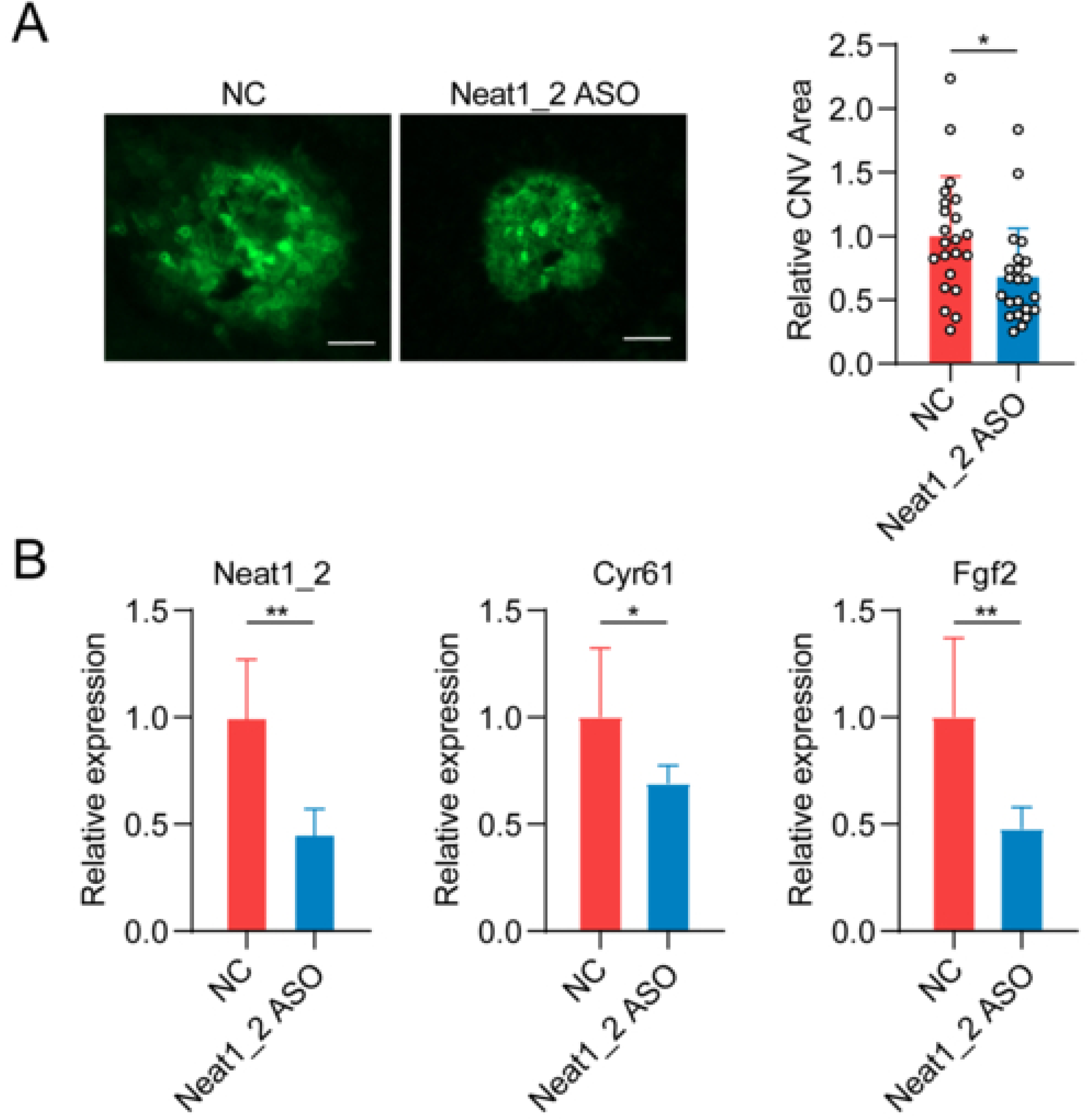
Intravitreal disruption of paraspeckle assembly using ASO targeting *NEAT1_2* Domain C suppresses neovascular remodeling and restores endothelial homeostasis in vivo. **(A)** Isolectin-B4-staining of choroidal flat mounts demonstrates significant suppression of CNV following intravitreal injection of an ASO targeting *NEAT1_2* Domain C. Scale bar = 100 μm. (**B**) qPCR analysis of Cyr61/Fgf2 expression in the mouse retinal (n = 6). Data are shown as mean ± SD from ≥ 3 independent repeats. \**p* < 0.05, ** *p* < 0.01, *** *p* < 0.001.

## Discussion

Our results showed a role of paraspeckles in mediating the relay from microbial insult-induced inflammation triggered by microbial insults into neovascular remodeling. Inflammation-induced *NEAT1_2* upregulation and paraspeckle hyper-assembly sequester RBM14 into paraspeckles and relieve its transcription repression of CYR61, which is amplified via CYR61 paracrine and FGF2 upregulation to drive neovascular remodeling. Our results show that paraspeckle dynamics act as regulatory switches in the link between microbial insults and host vascular endothelial homeostasis.

Recent studies have highlighted that paraspeckles, which are membrane-less nuclear bodies, play an important regulatory role in cellular responses to stress during tumorigenesis and viral infections ^17,34^. *NEAT1_2* serves as a structural scaffold for paraspeckle assembly, and its Domain C is a required structural component ^35–37^. Our results of ASO targeting Domain C and 1,6-HD treatment suggest that paraspeckle assembly plays a substantial role in governing vascular endothelial homeostasis. Moreover, our results further highlight the role of paraspeckle hyper-assembly in mediating the relay from microbial insults to compromized endothelial homeostasis and neovascular remodeling. Notably, our gut dysbiosis model results demonstrate that such a system-to-local relay is an inevitable component of the gut-brain/retinal axis, allowing microbial insults to influence the CNS ^38–40^. Our findings thus underscore paraspeckles as biophysical switches that translate microbial insult-induced inflammation into localized transcriptional programs.

Our findings highlighted the sequestration of RBM14 by paraspeckles in translating microbial insult-induced inflammation into gene expression response during neovascular remodeling. The RBM14 gene encodes an RNA-binding protein with multiple functions in transcriptional repression and splicing regulation ^41–43^, and maintains a basal “transcriptional brake” on the *CYR61* promoter under endothelial quiescent conditions. Such a spatial redistribution relieves transcriptional repression, triggering a cascade that increases CYR61 expression.

Furthermore, CYR61 enhanced FGF2 signaling in endothelial cells in a paracrine manner, highlighting the importance of CYR61-FGF2 signaling in endothelial homeostasis. CYR61 and FGF2 have been known as distinct signaling proteins that play complex roles in neurodegenerative diseases such as AD, AMD, and Parkinson’s disease. CYR61, also known as CCN1, is involved in cell adhesion, migration, proliferation, angiogenesis, and apoptosis. As a secreted extracellular matrix (ECM) protein, it plays a critical role in wound healing, skeletal development, and cardiovascular development. In addition, it physically displaces FGF2 from the extracellular matrix and heparin sulfate proteoglycans. By releasing FGF2 from its tethered state in the ECM, CYR61 increases local FGF2 bioavailability ^44–47^. Consistently, our results demonstrated that CYR61 upregulates FGF2 expression in a paracrine manner in endothelial cells, thereby sustaining a pathological remodeling loop.

Moreover, our intravitreal delivery of Domain C-targeted ASOs restores endothelial homeostasis and suppresses pathological remodeling in vivo. Given the conserved nature of the *NEAT1_2*/CYR61/FGF2 signature across human cohorts and mouse models as shown in our comparative transcriptome analysis, such findings thus highlight paraspeckles as a druggable target for endothelial homeostasis and an alternative to conventional anti-VEGF therapies ^48,49^.

There are also limitations in our current study. The laser-induced CNV used in our in vivo study is an acute inflammatory model, whereas chronic inflammation in aging and metabolic diseases warrants further study.

In summary, in our current study we demonstrate a substantial role of paraspeckles in mediating the systemic-to-local inflammatory relay during disrupted endothelial homeostasis and neurovascular remodeling by sequestering RBM14 and enhancing CYR61-FGF2 paracrine signaling. Furthermore, our study underscores paraspeckle assembly as a promising therapeutic target for neurovascular remodeling and related diseases.

## Acknowledgements

The graphical abstract was created using BioRender.com

## CRediT authorship contribution statement

J-QP, K-TY, and J-QZ performed the experiments and data collection. JQ-P and J-HC interpreted the data and performed statistical analysis. J-QP and J-QZ wrote the draft of the manuscript. Y-YJ and J-HC assisted in data analysis and critically revised the manuscript. All authors read and approved the final manuscript.

## Data availability

The 16S rRNA sequencing and RNA-Seq data used in this study have been deposited in the

Genome Sequence Archive at the National Genomics Data Center, Beijing Institute of Genomics, Chinese Academy of Sciences / China National Center for Bioinformation under the accession GSA: PRJCA018834, PRJCA018644, and GSA-human: PRJCA042371.

**Summary Graph. Diagram illustrating system-to-local inflammatory relay mediated by paraspeckle assembly during neurovascular remodeling.**

(1) Systemic inflammation: Gut dysbiosis or sepsis drives LPS influx, initiating a systemic-to-local inflammatory relay.

(2) Local decoding via RBM14 sequestration: *NEAT1_2* upregulation induced by systemic inflammation promotes paraspeckle hyper-assembly, which in turn sequesters RBM14, an RNA-binding protein functioning as a transcriptional brake on *CYR61* transcription.

(3) Paracrine Amplification: Spatial removal of RBM14 from the *CYR61* promoter relieves transcription repression, activating the CYR61/FGF2 signaling and neovascular remodeling. Secreted CYR61 mediates intercellular endothelial communication in a paracrine manner, amplifying CYR61/FGF2 signaling and driving localized neovascular remodeling.

(4) Therapeutic target: Disrupting paraspeckle assembly by targeting Domain C intercepts neurovascular remodeling and restores endothelial homeostasis.

Abbreviations: PAD, paraspeckle assembly domain. BBB/BRB, blood-brain and blood-retinal barriers.

## References

1. Chen AQ, Fang Z, Chen XL, Yang S, Zhou YF, Mao L, Xia YP, Jin HJ, Li YN, You MF, Wang XX, Lei H, He QW, Hu B. Microglia-derived TNF-α mediates endothelial necroptosis aggravating blood brain-barrier disruption after ischemic stroke. Cell Death Dis. 2019;10(7):487. doi:10.1038/s41419-019-1716-9

2. Zlokovic BV. Neurovascular pathways to neurodegeneration in Alzheimer’s disease and other disorders. Nat Rev Neurosci. 2011;12(12):723–738. doi:10.1038/nrn3114

3. Sekino N, Selim M, Shehadah A. Sepsis-associated brain injury: underlying mechanisms and potential therapeutic strategies for acute and long-term cognitive impairments. J Neuroinflammation. 2022;19(1):101. doi:10.1186/s12974-022-02464-4

4. Belkaid Y, Hand TW. Role of the microbiota in immunity and inflammation. Cell. 2014;157(1):121–141. doi:10.1016/j.cell.2014.03.011

5. Cauwels A, Rogge E, Vandendriessche B, Shiva S, Brouckaert P. Extracellular ATP drives systemic inflammation, tissue damage and mortality. Cell Death Dis. 2014;5(3):e1102–e1102. doi:10.1038/cddis.2014.70

6. Huang X, Qi J, Su Y, Zhou Y, Wang Q, Huang T, Xue D, Zeng Y, Verkhratsky A, Zhou B, Chen H, Yi C. Endothelial DR6 in blood-brain barrier malfunction in Alzheimer’s disease. Cell Death Dis. 2024;15(4):258. doi:10.1038/s41419-024-06639-0

7. Patton N, Pattie A, MacGillivray T, Aslam T, Dhillon B, Gow A, Starr JM, Whalley LJ, Deary IJ. The association between retinal vascular network geometry and cognitive ability in an elderly population. Invest Ophthalmol Vis Sci. 2007;48(5):1995–2000. doi:10.1167/iovs.06-1123

8. Patton N, Aslam T, Macgillivray T, Pattie A, Deary IJ, Dhillon B. Retinal vascular image analysis as a potential screening tool for cerebrovascular disease: a rationale based on homology between cerebral and retinal microvasculatures. J Anat. 2005;206(4):319–348. doi:10.1111/j.1469-7580.2005.00395.x

9. Sweeney MD, Zhao Z, Montagne A, Nelson AR, Zlokovic BV. Blood-Brain Barrier: From Physiology to Disease and Back. Physiol Rev. 2019;99(1):21–78. doi:10.1152/physrev.00050.2017

10. Ting DSW, Pasquale LR, Peng L, Campbell JP, Lee AY, Raman R, Tan GSW, Schmetterer L, Keane PA, Wong TY. Artificial intelligence and deep learning in ophthalmology. Br J Ophthalmol. 2019;103(2):167–175. doi:10.1136/bjophthalmol-2018-313173

11. Hirose T, Yamazaki T, Nakagawa S. Molecular anatomy of the architectural NEAT1 noncoding RNA: The domains, interactors, and biogenesis pathway required to build phase-separated nuclear paraspeckles. Wiley Interdiscip Rev RNA. 2019;10(6):e1545. doi:10.1002/wrna.1545

12. Li M, Li M, Xia Y, Li G, Su X, Wang D, Ye J, Lu F, Sun T, Ji C. HDAC1/3-dependent moderate liquid-liquid phase separation of YY1 promotes METTL3 expression and AML cell proliferation. Cell Death Dis. 2022;13(11):992. doi:10.1038/s41419-022-05435-y

13. Watanabe S, Inami H, Oiwa K, Murata Y, Sakai S, Komine O, Sobue A, Iguchi Y, Katsuno M, Yamanaka K. Aggresome formation and liquid-liquid phase separation independently induce cytoplasmic aggregation of TAR DNA-binding protein 43. Cell Death Dis. 2020;11(10):909. doi:10.1038/s41419-020-03116-2

14. Zhang Y, Luo M, Cui X, O’Connell D, Yang Y. Long noncoding RNA NEAT1 promotes ferroptosis by modulating the miR-362-3p/MIOX axis as a ceRNA. Cell Death Differ. 2022;29(9):1850–1863. doi:10.1038/s41418-022-00970-9

15. Fox AH, Lamond AI. Paraspeckles. Cold Spring Harb Perspect Biol. 2010;2(7):a000687. doi:10.1101/cshperspect.a000687

16. Bond CS, Fox AH. Paraspeckles: nuclear bodies built on long noncoding RNA. J Cell Biol. 2009;186(5):637–644. doi:10.1083/jcb.200906113

17. Patil A, Strom AR, Paulo JA, Collings CK, Ruff KM, Shinn MK, Sankar A, Cervantes KS, Wauer T, St Laurent JD, Xu G, Becker LA, Gygi SP, Pappu RV, Brangwynne CP, Kadoch C. A disordered region controls cBAF activity via condensation and partner recruitment. Cell. 2023;186(22):4936–4955.e26. doi:10.1016/j.cell.2023.08.032

18. Liu YS, Pan JQ, Pan XB, Kong FS, Zhang JQ, Wei ZY, Xu ZH, Rao JH, Wang JH, Chen JH. Comparative Analysis of Molecular Landscape in Mouse Models and Patients Reveals Conserved Inflammation Pathways in Age-Related Macular Degeneration. Invest Ophthalmol Vis Sci. 2024;65(1):13. doi:10.1167/iovs.65.1.13

19. Bolyen E, Rideout JR, Dillon MR, Bokulich NA, Abnet CC, Al-Ghalith GA, Alexander H, Alm EJ, Arumugam M, Asnicar F, Bai Y, Bisanz JE, Bittinger K, Brejnrod A, Brislawn CJ, Brown CT, Callahan BJ, Caraballo-Rodríguez AM, Chase J, Cope EK, Da Silva R, Diener C, Dorrestein PC, Douglas GM, Durall DM, Duvallet C, Edwardson CF, Ernst M, Estaki M, Fouquier J, Gauglitz JM, Gibbons SM, Gibson DL, Gonzalez A, Gorlick K, Guo J, Hillmann B, Holmes S, Holste H, Huttenhower C, Huttley GA, Janssen S, Jarmusch AK, Jiang L, Kaehler BD, Kang KB, Keefe CR, Keim P, Kelley ST, Knights D, Koester I, Kosciolek T, Kreps J, Langille MGI, Lee J, Ley R, Liu YX, Loftfield E, Lozupone C, Maher M, Marotz C, Martin BD, McDonald D, McIver LJ, Melnik AV, Metcalf JL, Morgan SC, Morton JT, Naimey AT, Navas-Molina JA, Nothias LF, Orchanian SB, Pearson T, Peoples SL, Petras D, Preuss ML, Pruesse E, Rasmussen LB, Rivers A, Robeson MS, Rosenthal P, Segata N, Shaffer M, Shiffer A, Sinha R, Song SJ, Spear JR, Swafford AD, Thompson LR, Torres PJ, Trinh P, Tripathi A, Turnbaugh PJ, Ul-Hasan S, van der Hooft JJJ, Vargas F, Vázquez-Baeza Y, Vogtmann E, von Hippel M, Walters W, Wan Y, Wang M, Warren J, Weber KC, Williamson CHD, Willis AD, Xu ZZ, Zaneveld JR, Zhang Y, Zhu Q, Knight R, Caporaso JG. Reproducible, interactive, scalable and extensible microbiome data science using QIIME 2. Nat Biotechnol. 2019;37(8):852–857. doi:10.1038/s41587-019-0209-9

20. Segata N, Izard J, Waldron L, Gevers D, Miropolsky L, Garrett WS, Huttenhower C. Metagenomic biomarker discovery and explanation. Genome Biol. 2011;12(6):R60. doi:10.1186/gb-2011-12-6-r60

21. Ratnapriya R, Sosina OA, Starostik MR, Kwicklis M, Kapphahn RJ, Fritsche LG, Walton A, Arvanitis M, Gieser L, Pietraszkiewicz A, Montezuma SR, Chew EY, Battle A, Abecasis GR, Ferrington DA, Chatterjee N, Swaroop A. Retinal transcriptome and eQTL analyses identify genes associated with age-related macular degeneration. Nat Genet. 2019;51(4):606–610. doi:10.1038/s41588-019-0351-9

22. Griggs E, Trageser K, Naughton S, Yang EJ, Mathew B, Van Hyfte G, Hellmers L, Jette N, Estill M, Shen L, Fischer T, Pasinetti GM. Recapitulation of pathophysiological features of AD in SARS-CoV-2-infected subjects. eLife. 2023;12:e86333. doi:10.7554/eLife.86333

23. Pinheiro da Silva F, Gonçalves ANA, Duarte-Neto AN, Dias TL, Barbeiro HV, Breda CNS, Breda LCD, Câmara NOS, Nakaya HI. Transcriptome analysis of six tissues obtained post-mortem from sepsis patients. J Cell Mol Med. 2023;27(20):3157–3167. doi:10.1111/jcmm.17938

24. Orozco LD, Owen LA, Hofmann J, Stockwell AD, Tao J, Haller S, Mukundan VT, Clarke C, Lund J, Sridhar A, Mayba O, Barr JL, Zavala RA, Graves EC, Zhang C, Husami N, Finley R, Au E, Lillvis JH, Farkas MH, Shakoor A, Sherva R, Kim IK, Kaminker JS, Townsend MJ, Farrer LA, Yaspan BL, Chen HH, DeAngelis MM. A systems biology approach uncovers novel disease mechanisms in age-related macular degeneration. Cell Genomics. 2023;3(6):100302. doi:10.1016/j.xgen.2023.100302

25. Chen S, Zhou Y, Chen Y, Gu J. fastp: an ultra-fast all-in-one FASTQ preprocessor. Bioinforma Oxf Engl. 2018;34(17):i884–i890. doi:10.1093/bioinformatics/bty560

26. Dobin A, Gingeras TR. Mapping RNA-seq Reads with STAR. Curr Protoc Bioinforma. 2015;51:11.14.1–11.14.19. doi:10.1002/0471250953.bi1114s51

27. Li H, Handsaker B, Wysoker A, Fennell T, Ruan J, Homer N, Marth G, Abecasis G, Durbin R, 1000 Genome Project Data Processing Subgroup. The Sequence Alignment/Map format and SAMtools. Bioinforma Oxf Engl. 2009;25(16):2078–2079. doi:10.1093/bioinformatics/btp352

28. Robinson MD, McCarthy DJ, Smyth GK. edgeR: a Bioconductor package for differential expression analysis of digital gene expression data. Bioinforma Oxf Engl. 2010;26(1):139–140. doi:10.1093/bioinformatics/btp616

29. Kuleshov MV, Jones MR, Rouillard AD, Fernandez NF, Duan Q, Wang Z, Koplev S, Jenkins SL, Jagodnik KM, Lachmann A, McDermott MG, Monteiro CD, Gundersen GW, Ma’ayan A. Enrichr: a comprehensive gene set enrichment analysis web server 2016 update. Nucleic Acids Res. 2016;44(W1):W90–97. doi:10.1093/nar/gkw377

30. Huang DW, Sherman BT, Lempicki RA. Bioinformatics enrichment tools: paths toward the comprehensive functional analysis of large gene lists. Nucleic Acids Res. 2009;37(1):1–13. doi:10.1093/nar/gkn923

31. Pertea M, Pertea GM, Antonescu CM, Chang TC, Mendell JT, Salzberg SL. StringTie enables improved reconstruction of a transcriptome from RNA-seq reads. Nat Biotechnol. 2015;33(3):290–295. doi:10.1038/nbt.3122

32. Love MI, Huber W, Anders S. Moderated estimation of fold change and dispersion for RNA-seq data with DESeq2. Genome Biol. 2014;15(12):550. doi:10.1186/s13059-014-0550-8

33. Yamazaki T, Souquere S, Chujo T, Kobelke S, Chong YS, Fox AH, Bond CS, Nakagawa S, Pierron G, Hirose T. Functional domains of NEAT1 architectural lncRNA induce paraspeckle assembly through phase separation. Mol Cell. 2018;70(6):1038–1053.e7. doi:10.1016/j.molcel.2018.05.019

34. Ku D, Yang Y, Kim Y. RNA-associated nuclear condensates: Where the nucleus keeps its RNAs in check. Mol Cells. 2025;48(8):100240. doi:10.1016/j.mocell.2025.100240

35. Clemson CM, Hutchinson JN, Sara SA, Ensminger AW, Fox AH, Chess A, Lawrence JB. An Architectural Role for a Nuclear Noncoding RNA: NEAT1 RNA Is Essential for the Structure of Paraspeckles. Mol Cell. 2009;33(6):717–726. doi:10.1016/j.molcel.2009.01.026

36. Banani SF, Lee HO, Hyman AA, Rosen MK. Biomolecular condensates: organizers of cellular biochemistry. Nat Rev Mol Cell Biol. 2017;18(5):285–298. doi:10.1038/nrm.2017.7

37. Mao YS, Sunwoo H, Zhang B, Spector DL. Direct visualization of the co-transcriptional assembly of a nuclear body by noncoding RNAs. Nat Cell Biol. 2011;13(1):95–101. doi:10.1038/ncb2140

38. Andriessen EM, Wilson AM, Mawambo G, Dejda A, Miloudi K, Sennlaub F, Sapieha P. Gut microbiota influences pathological angiogenesis in obesity-driven choroidal neovascularization. EMBO Mol Med. 2016;8(12):1366–1379. doi:10.15252/emmm.201606531

39. Rowan S, Jiang S, Korem T, Szymanski J, Chang ML, Szelog J, Cassalman C, Dasuri K, McGuire C, Nagai R, Du XL, Brownlee M, Rabbani N, Thornalley PJ, Baleja JD, Deik AA, Pierce KA, Scott JM, Clish CB, Smith DE, Weinberger A, Avnit-Sagi T, Lotan-Pompan M, Segal E, Taylor A. Involvement of a gut–retina axis in protection against dietary glycemia-induced age-related macular degeneration. Proc Natl Acad Sci. 2017;114(22):E4472–E4481. doi:10.1073/pnas.1702302114

40. Rowan S, Taylor A. Gut microbiota modify risk for dietary glycemia-induced age-related macular degeneration. Gut Microbes. 2018;9(5):452–457. doi:10.1080/19490976.2018.1435247

41. Hennig S, Kong G, Mannen T, Sadowska A, Kobelke S, Blythe A, Knott GJ, Iyer KS, Ho D, Newcombe EA, Hosoki K, Goshima N, Kawaguchi T, Hatters D, Trinkle-Mulcahy L, Hirose T, Bond CS, Fox AH. Prion-like domains in RNA binding proteins are essential for building subnuclear paraspeckles. J Cell Biol. 2015;210(4):529–539. doi:10.1083/jcb.201504117

42. Li J, Wang C, Feng G, Zhang L, Chen G, Sun H, Wang J, Zhang Y, Zhou Q, Li W. Rbm14 maintains the integrity of genomic DNA during early mouse embryogenesis via mediating alternative splicing. Cell Prolif. 2020;53(1):e12724. doi:10.1111/cpr.12724

43. Li J, Wang Y, Rao X, Wang Y, Feng W, Liang H, Liu Y. Roles of alternative splicing in modulating transcriptional regulation. BMC Syst Biol. 2017;11(5):89. doi:10.1186/s12918-017-0465-6

44. Menéndez JA, Mehmi I, Griggs DW, Lupu R. The angiogenic factor CYR61 in breast cancer: molecular pathology and therapeutic perspectives. Endocr Relat Cancer. 2003;10(2):141–152. doi:10.1677/erc.0.0100141

45. You JJ, Yang CH, Chen MS, Yang CM. Cysteine-rich 61, a Member of the CCN Family, as a Factor Involved in the Pathogenesis of Proliferative Diabetic Retinopathy. Invest Ophthalmol Vis Sci. 2009;50(7):3447–3455. doi:10.1167/iovs.08-2603

46. Schmitz P, Gerber U, Schütze N, Jüngel E, Blaheta R, Naggi A, Torri G, Bendas G. Cyr61 is a target for heparin in reducing MV3 melanoma cell adhesion and migration via the integrin VLA-4. Thromb Haemost. 2013;110(5):1046–1054. doi:10.1160/TH13-02-0158

47. Marinkovic M, Dai Q, Gonzalez AO, Tran ON, Block TJ, Harris SE, Salmon AB, Yeh CK, Dean DD, Chen XD. Matrix-bound Cyr61/CCN1 is required to retain the properties of the bone marrow mesenchymal stem cell niche but is depleted with aging. Matrix Biol. 2022;111:108–132. doi:10.1016/j.matbio.2022.06.004

48. Xu M, Fan R, Fan X, Shao Y, Li X. Progress and challenges of anti-VEGF agents and their sustained-release strategies for retinal angiogenesis. Drug Des Devel Ther. 2022;16:3241–3262. doi:10.2147/DDDT.S383101

49. Dinah C, Dodds M, Lotery A, Salvatore S, Fletcher E, Lake AVR, Parker A, Paris Pereira L, Retiere AC, Saffar I, Arrisi P, Chi GC. Treatment patterns and long-term outcomes in anti-VEGF-treated macular oedema secondary to retinal vein occlusion: A retrospective observational study. Eye. 2026;40(1):107–116. doi:10.1038/s41433-025-04089-2

